# Catabolism of alkylphenols in *Rhodococcus* via a *meta*-cleavage pathway associated with genomic islands

**DOI:** 10.1101/674713

**Authors:** David J. Levy-Booth, Morgan M. Fetherolf, Gordon Stewart, Jie Liu, Lindsay D. Eltis, William W. Mohn

## Abstract

The bacterial catabolism of aromatic compounds has considerable promise to convert lignin depolymerization products to commercial chemicals. Alkylphenols are a key class of depolymerization products whose catabolism is not well elucidated. We isolated *Rhodococcus rhodochrous* EP4 on 4-ethylphenol and applied genomic and transcriptomic approaches to elucidate alkylphenol catabolism in EP4 and *Rhodococcus jostii* RHA1. RNA-Seq and RT-qPCR revealed a pathway encoded by the *aphABCDEFGHIQRS* genes that degrades 4-ethylphenol via the *meta-*cleavage of 4-ethylcatechol. This process was initiated by a two-component alkylphenol hydroxylase, encoded by the *aphAB* genes, which were up-regulated ~3,000-fold. Purified AphAB from EP4 had highest specific activity for 4-ethylphenol and 4-propylphenol (~2000 U/mg) but did not detectably transform phenol. Nevertheless, a Δ*aphA* mutant in RHA1 grew on 4-ethylphenol by compensatory up-regulation of phenol hydroxylase genes (*pheA1-3*). Deletion of *aphC*, encoding an extradiol dioxygenase, prevented growth on 4-alkylphenols but not phenol. Disruption of *pcaL* in the β-ketoadipate pathway prevented growth on phenol but not 4-alkylphenols. Thus, 4-ethylphenol and 4-propylphenol are catabolized exclusively via *meta-*cleavage in rhodococci while phenol is subject to *ortho-*cleavage. Putative genomic islands encoding *aph* gene*s* were identified in EP4 and several other rhodococci. Overall, this study identifies a 4-alkylphenol pathway in rhodococci, demonstrates key enzymes involved, and presents evidence that the pathway is encoded in a genomic island. These advances are of particular importance for wide-ranging industrial applications of rhodococci, including upgrading of lignocellulose biomass.

**Importance:** Elucidation of bacterial alkylphenol catabolism is important for the development of biotechnologies to upgrade the lignin component of plant biomass. We isolated a new strain, *Rhodococcus rhodochrous* EP4, on 4-ethylphenol, an alkylphenol that occurs in lignin-derived streams, including reductive catalytic fractionation products of corn stover. We further demonstrated its degradation via a *meta-*cleavage pathway (Aph) with transcriptomics. A new class of Actinobacterial hydroxylase, AphAB, acts specifically on alkylphenols. Phylogenomic analysis indicated that the *aph* genes occur on putative genomic islands in several rhodococcal strains. These genes were identified in the genetically-tractable strain *Rhodococcus jostii* RHA1. Strains missing this element cannot metabolise 4-ethylphenol and 4-propylphenol. Overall, we advanced the understanding of how aromatic compounds are degraded by environmental bacteria and identified enzymes that can be employed in the transition away from petro-chemicals towards renewable alternatives.

## Introduction

Lignin, a heterogeneous aromatic polymer, accounts for up to 40% dry weight of terrestrial plant biomass (Ragauskas et al., 2014). It is primarily composed of *p*-hydroxyphenyl (H), guaiacyl (G), and sinapyl (S) subunits, polymerized by ether and C-C bonds (Boerjan et al., 2003). Lignin’s heterogeneity and recalcitrant bonds create substantial barriers to its efficient microbial and chemical degradation. Industrial lignin depolymerization is gaining traction as a means to produce fuels and chemicals historically derived from petroleum (Beckham et al., 2016; Ragauskas et al., 2014). Yet heterogeneous depolymerization products can require extensive separation and purification (Linger et al., 2014; Ragauskas et al., 2014). Bacterial biocatalysts provide a means of transforming mixtures of aromatic compounds to single compounds due to the convergent nature of their catabolic pathways, whereby substrates are transformed to central metabolites via shared intermediates, such as catechols (Eltis and Singh, 2018; Linger et al., 2014). Harnessing this biological funneling to refine lignin to high-value chemicals (Beckham et al., 2016; Eltis and Singh, 2018; Linger et al., 2014) is limited in part by a lack of knowledge of the catabolism of lignin-derived monomers.

Alkylphenols are a major class of aromatic compounds generated by a variety of lignin depolymerization technologies. For example, solvolysis of corn lignin produced 24 wt % alkylated monolignins, 46% of which was 4-ethylphenol derived from H-subunits (Jiang et al., 2014). Alkylphenols were also major pyrolysis products of wheat straw black liquor lignin fractions (Guo et al., 2017). Existing depolymerization strategies can require multiple stages of preprocessing and depolymerization, high heat or corrosive chemicals, and can produce dozens of alkylphenol and aromatic products (Asawaworarit et al., 2019; Kim et al., 2015; Ye et al., 2012). One promising depolymerization strategy that produces a narrow stream of alkylphenols is reductive catalytic reduction (RCF) (Pepper and Lee, 1969). 4-Ethylphenol was a major RCF product of corn stover, comprising up to 16.4% of the resulting aromatic monomers (Anderson et al., 2016).

Two bacterial pathways for the aerobic catabolism of 4-ethylphenol have been reported, initially involving either oxidation of the alkyl-side chain or hydroxylation of the aromatic ring. In *Pseudomonas putida* JD1, the alkyl-side chain is oxidized by 4-ethylphenol methylhydroxylase to eventually yield hydroquinone (Darby et al., 1987; Hopper and Cottrell, 2003). In contrast, *Pseudomonas* sp. KL28 hydroxylates 4-ethylphenol to 4-ethylcatechol (Jeong et al., 2003). In these pathways, the hydroquinone and 4-ethylcatechol undergo *meta*-cleavage (Darby et al., 1987; Jeong et al., 2003). In KL28, the alkylphenol hydroxylase is a six-component enzyme encoded by genes organized in a single co-linear transcriptional unit within a *meta*-cleavage pathway gene cluster (Jeong et al., 2003). A homologous pathway degrades phenol in *Comamonas testosteroni* TA441 (Arai et al., 2000).

*Rhodococcus* is a genus of mycolic acid-producing Actinobacteria that catabolize a wide variety of aromatic compounds (Yam et al., 2010), including phenols (Gröning et al., 2014; Kolomytseva et al., 2007). These bacteria also have considerable potential as biocatalysts for the industrial production of compounds ranging from nitriles, to steroids and high-value lipids (Alvarez et al., 1996; Round et al., 2017; Sengupta et al., 2019; Shields-Menard et al., 2017). In *Rhodococcus*, phenol catabolism is initiated by a two-component flavin-dependent monooxygenase (PheA1A2) (Saa et al. 2010) to generate a catechol. PheA1A2 homologs in *Rhodococcus opacus* 1CP can also hydroxylate chlorophenols and 4-methylphenol (Gröning et al., 2014) to produce the corresponding catechols, which undergo *ortho-*cleavage (Kolomytseva et al., 2007; Maltseva et al., 1994) and subsequent transformation to central metabolites via the β-ketoadipate pathway. In *rhodococci*, the β-ketoadipate pathway is confluent, with branches responsible for protocatechuate and catechol catabolism converging at PcaL, a β-ketoadipate enol-lactonase (Patrauchan et al., 2005; Yam et al., 2010). However, some *rhodococci* appear to have pathways responsible for the catabolism of alkylated aromatic compounds via *meta*-cleavage (Jang et al., 2005). Elucidating 4-alkylphenol metabolism in *rhodococci* will improve our understanding of Actinobacterial aromatic degradation and support the development of *Rhodococcus* strains as platforms for industrial lignin upgrading.

Genomic islands (GIs) are DNA segments likely to have been acquired by horizontal gene transfer. They are characterized by altered nucleotide characteristics (e.g., GC content), syntenic conservation, and frequent presence of mobility genes (transposases, insertion sequences (IS), and integrases) (Hacker and Kaper, 2000; Juhas et al., 2009). They can be further identified by the absence of genomic regions in closely-related strains (Hacker et al., 1990). GIs can confer resistance, virulence, symbiosis, and catabolic pathways (Dobrindt et al., 2004; Juhas et al., 2009). For example, the self-transferable *clc* element enabling 3- and 4-chlorocatechol and 2-aminophenol catabolism was identified as a GI in several *Gamma*- and *Betaproteobacteria* strains (Gaillard et al., 2006). Recent horizontal gene transfer may have played less of a role in shaping the *Rhodococcus jostii* RHA1 genome than in other bacteria such as *Burkholderia xenovorans* LB400, which has a similarly-sized genome (McLeod et al., 2006). Further, although RHA1 contains a high number of aromatic pathways, genes encoding these pathways are slightly underrepresented in the identified genomic islands. GIs can ameliorate in host genomes through nucleotide optimization or loss of mobility elements (Juhas et al., 2009; Lawrence and Ochman, 1997), reducing our effectiveness at predicting ancestral genomic additions. However, examination of GIs in multiple related genomes with an ensemble of predictive software can improve our understanding of the role of GIs in the evolution of bacterial catabolic pathways.

This study sought to identify catabolic pathways required for 4-alkylphenol catabolism. We report on the genomic, transcriptomic and enzymatic characterization of 4-alkylphenol catabolism in a newly-isolated 4-ethylphenol-degrading bacterium, *Rhodococcus rhodochrous* EP4 (Figure 1A), as well as RHA1. The activity of a novel two-component alkylphenol monooxygenase (AphAB) was characterized. Gene deletion analysis was employed to identify the subsequent route of catechol catabolism. Genomic analysis identified a putative *aph* GI, providing new evolutionary insight to the *aph meta-*cleavage pathway in *Rhodococcus*. Knowledge gained by this study will facilitate the efficient valorization of lignin following its depolymerization.

**Figure 1.**
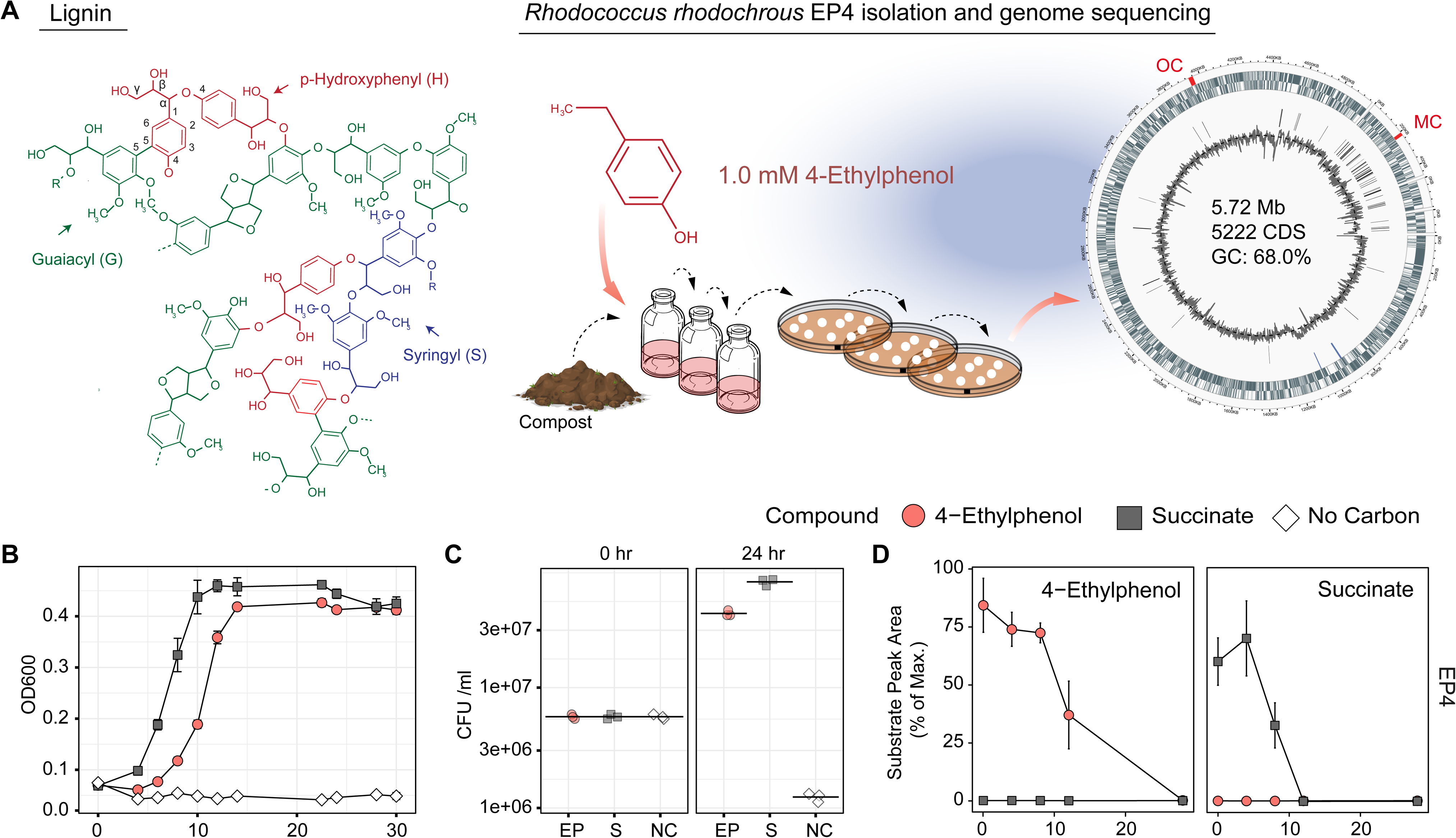
Isolation and growth of 4-ethylphenol-catabolizing strain *Rhodococcus rhodochrous* EP4. A) Schematic of EP4 isolation from compost on 4-ethylphenol, which is produced from reductive lignin depolymerization. Outer genomic track: coding sequences by strand; Inner track: insertion sequences; lines: GC content (%). MC, *meta*-cleavage pathway gene cluster; OC, *ortho*-cleavage pathway gene cluster. B) Growth of EP4 on 1 mM 4-ethylphenol or 2 mM succinate controls as optical density at 600 nm (OD_600_). Points and error bars reflect mean and standard error (n = 3). C) Colony forming units (CFU) during growth in Panel B. Points and horizontal bar indicate individual measurements and mean. D) Removal of 4-ethylphenol and succinate in EP4 cultures by GC/MS.

## Materials and Methods

### Bacterial strains and growth conditions

Liquid enrichment cultures were inoculated with either ~4 month aged agricultural compost (~25 cm depth) from The University of British Columbia farm (49° 14′ 57.8904″ N, 123° 14′ 0.0492″ W) or forest soil from Pacific Spirit Park in Vancouver, Canada. The cultures contained 1.0 mM 4-ethylphenol (≥97.0% Sigma-Aldrich, St. Louis, U.S.A.) as sole organic substrate in M9-Goodies (Bauchop and Elsden, 1960; Elder, 1983). The cultures were incubated at 30°C with shaking at 200 rpm for 2 weeks. Removal of 4-ethylphenol was monitored by GC-MS, after which cultures were transferred to fresh medium, 0.5% inocula. After 3 serial transfers, isolates were obtained by plating on homologous medium solidified with 1.5% purified agar. Colonies appeared in 10 days. Individual colonies were transferred to liquid media and replated on solid media for colony isolation. *Rhodococcus rhodochrous* strain DSM43241 was purchased from DSMZ (Braunschweig, Germany).

EP4 and RHA1 cultures for RNA extraction were grown overnight at 30°C on LB broth (200 rpm), diluted to 0.05 OD_600_ and washed thrice in M9 with no supplement, and grown to mid-log phase on 50 ml M9-Goodies with 1.0 mM 4-ethylphenol or succinate. EP4 was additionally grown on 1.0 mM benzoate (99%, Sigma-Aldrich). Additional screening of EP4, *R*. *rhodochrous* DSM43241, RHA1, and mutant strains used 5 ml M9-Goodies with 1.0 mM phenol (≥99%, VWR International, Ltd., Mississauga, Canada), 3-methylphenol (*m-*cresol; 99% Sigma-Aldrich), 4-methylphenol (99% Sigma-Aldrich), 4-propylphenol (>99%, TCI), 3,4-dimethylphenol (DMP) (99% Sigma-Aldrich), 2,4-DMP (98% Sigma-Aldrich), 4-hydroxyphenylacetate (HPA) (98% Sigma-Aldrich), 4-hydroxybenzioic acid (HBA) (99% Sigma-Aldrich), or 0.5 mM 4-nitrophenol (NP) (≥99%, Sigma-Aldrich), incubated for 24 h. Cells were lysed by boiling in 1M NaOH and protein quantified using the Micro BCA™ Protein Assay (Thermo-Fisher Scientific Inc., Waltham, U.S.A.) and a VersaMax™ microplate reader (Molecular Devices LLC, San Jose, U.S.A.).

### DNA manipulation, plasmid construction and gene deletion

DNA was isolated, manipulated, and analyzed using standard protocols (Sambrock and W. Russel, 2001). *E. coli* and RHA1 were transformed with DNA by electroporation using a MicroPulser with GenePulser cuvettes (Bio-Rad). To produce N-terminal His_6_-tagged Aph_AEP4_, AphB_EP4_ and AphC_RHA1_ (See Table 1) C6369_RS01585, C6369_RS01550 and RHA1_RS18750 were amplified from genomic DNA using Phusion Polymerase™ with the oligonucleotides listed in Supplementary Table 1. The nucleotide sequence of the cloned genes was verified. The *ΔaphA* and *ΔaphC* mutants were constructed using a *sacB* counter selection system (van der Geize et al., 2007). Five hundred bp flanking regions of RHA1_RS18785 and RHA1_RS18750 were amplified from RHA1 genomic DNA using the primers listed in Supplementary Table 1. The resulting amplicons were inserted into pK18mobsacB linearized with EcoR1 using Gibson Assembly. The nucleotide sequences of the resulting constructs were verified. Kanamycin-sensitive/sucrose-resistant colonies were screened using PCR and the gene deletion was confirmed by sequencing.

**Table 1.**
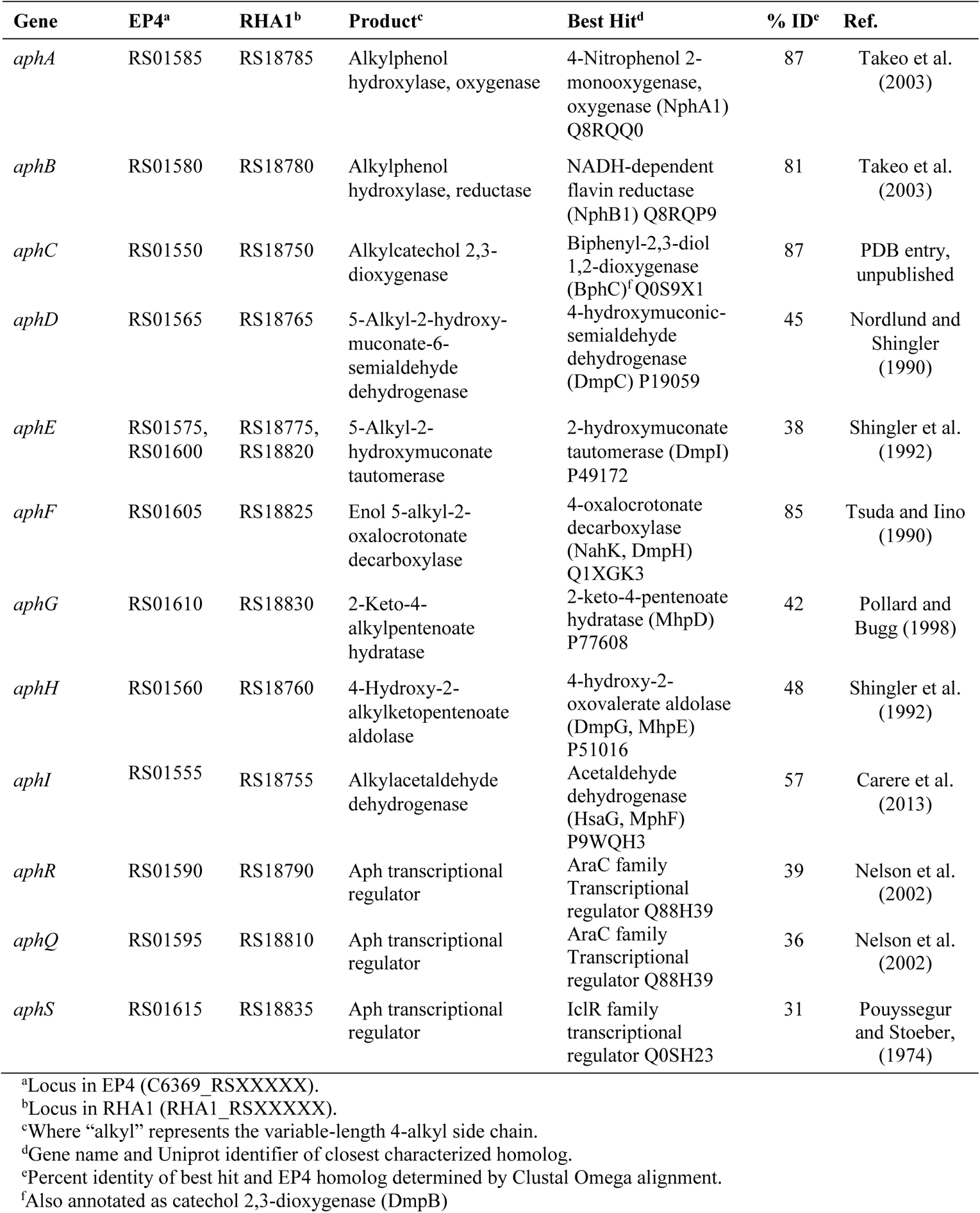
Genes in the alkylphenol *meta*-cleavage pathway

### Enzyme production and purification

The production and purification of AphA_EP4_, AphB _EP4_ and AphC_RHA1_ are described in Supplementary Methods.

### GC/MS analysis

Growth substrate depletion was analyzed in culture supernatants using an Agilent Technologies (Santa Clara, U.S.A.) 6890N gas chromatograph equipped with a 30-m Agilent 190915-433 capillary column and a 5973 mass-selective detector (GC/MS). Briefly, 400-µl samples of culture supernatant were amended with 3-chlorobenzoic acid (as internal standard), extracted with ethyl acetate (1:1 v/v) and dried under a nitrogen stream. The extract was suspended in pyridine and derivatized with trimethylsilyl for 1 h at 60°C. One-µl injections were analyzed using the following parameters: transfer line temperature of 325°C, run temperature of 90°C for 3 min, then ramped to 290°C at 12°C min^−1^ with a 10 min final hold. Peaks from raw trace files were aligned and integrated using *xcms* in R 3.4.4 (R Core Team, 2016) against 4-ethylphenol and succinate standards. Values were normalized to the area of the internal standard and expressed as a percent of maximum peak area.

### Nucleic acid extraction and sequencing

RNA was extracted from cellular pellets from 15 ml of EP4 and RHA1 cultures using TRIzol™ (Thermo-Fisher) and Turbo™ DNase (Thermo-Fisher). Quality and quantity of nucleic acids were assessed using 1% [w/v] agarose gel electrophoresis and Qubit fluorometric quantitation (Thermo-Fisher), prior to storage at −80°C. Approximately 1 µg RNA underwent Ribo-Zero rRNA removal (Illumina, San Diego, U.S.A.), TruSeq LT (Illumina) library preparation, and sequencing using HiSeq4000 2×100bp. Genomic DNA was extracted using CTAB. Fifteen µg was pulse-field electrophoresis size-selected and sequenced with one Pacific Biosciences (PacBio) RS II SMRT cell.

### Bioinformatics

A *de novo* draft genome was assembled with HGAP in the SMRT Analysis v2.3 pipeline (Chin et al., 2013) and MeDuSa 1.6 scaffolding (Bosi et al., 2015). Circularizing the genome sequence was attempted using Circlator 1.5.5 (Hunt et al., 2015), and plasmid detection was attempted using PlasmidFinder 2.0.1 (Carattoli et al., 2014). Annotation used the NCBI Prokaryotic Genome Annotation Pipeline (PGAP) 4.4 and BLASTp against the Protein Data Bank (e-value 10^−3^). Quality control, filtering and trimming of RNA reads used Trimmomatic 0.3.6 defaults (Bolger et al., 2014). Assembly used Trinity 2.4.0 (Grabherr et al., 2011). Transcript quantification used HTSeq 0.9.1 (Anders et al., 2015), FeatureCounts 1.5.0-p3 (Liao et al., 2014) and Salmon 0.8.1 (Patro et al., 2017). Differentially-expression was analyzed using *DeSeq2* 1.18.1 (Love et al., 2014) with false discovery rate (*fdr*) correction. RT-qPCR conditions are in in Supplementary Methods. All data visualization used *ggplot2* 3.1.0 unless otherwise noted. To download all reference data and re-create the transcriptomic analysis herein, refer to: https://github.com/levybooth/Rhodococcus_Transcriptomics.

### Phylogenomic characterization

Protein sequences were structurally aligned with T-Coffee 11.00 Expresso (Armougom et al., 2006); maximum-likelihood trees were generated using the best of 100 RAxML 8.0.0 iterations using the PROTGAMMALG model (Stamatakis, 2014) and visualized with iTOL v4 (Letunic and Bork, 2016). Genomic island regions were predicted with IslandViewer 4 (Bertelli et al., 2017; Hsiao et al., 2003). All available *Rhodococcus* genomes (n = 325) were downloaded from NCBI RefSeq. Genomic alignment with a subset of genomes (Supplementary Table 2) used *nucmer* 3.1 (Marçais et al., 2018) and visualized with *circlize* 0.4.5 (Gu et al., 2014). Ribosomal protein trees were constructed as in Hug et al. (2016).

### Enzyme assays

AphAB_EP4_ activity was measured spectrophotometrically by following the *meta*-cleavage of the produced 4-ethylcatechol in a coupled assay with AphC_RHA1_ at 25 ± 0.5 °C. The standard assay was performed in 200μl air- saturated 20 mM MOPS, 90 mM NaCl (*I* = 0.1 mM, pH 7.2) containing 0.5 mM 4-ethylphenol, 5 μM AphA_EP4_, 1 μM AphB_EP4_, 0.2 μMAphC_RHA1_, 1000 U mL^−1^ of catalase, 1 mM NADH, and 2.5 μM FAD. Components were incubated for 30 s, then the reaction was initiated by adding NADH and was monitored at 400 nm. Absorbance was monitored using a Varian Cary 5000 spectrophotometer controlled by WinUV software. One unit of activity, U, was defined as the amount of enzyme required to hydroxylate of 1 nmol substrate per minute. Extinction coefficients for methyl-, ethyl-, and propylcatechol cleavage at 400 nm were 18,600, 15,100, 19,400 M^−1^ cm^−1^, respectively, calculated by differences in liberation of O2 from alkylcatechol cleavage by 0.2 nmol AphC_RHA1_ monitored using a Clark- type polarographic O_2_ electrode OXYG1 (Hansatech, Pentney, UK) connected to a circulating water bath. Details of additional enzyme end point assays are provided in the Supplementary Methods.

## Results

### Isolation and genomic characterization of a 4-ethylphenol-degrading *Rhodococcus* strain

Enrichment cultures with 4-ethylphenol as a sole organic growth substrate were inoculated with either forest soil or compost and incubated at 30°C. Those cultures inoculated with compost demonstrated superior potential for 4-ethylphenol degradation and were used for subsequent isolation of strain EP4. The 16S rRNA gene (27F-1492R; Lane, 1991) of EP4 shared 100% sequence identity with that of *R. rhodochrous* NBRC 16069. *De novo* assembly produced a 5.72-Mb, high-quality, single-scaffold EP4 genome sequence (Figure 1A) containing 5,198 predicted genes: 4,942 protein coding sequences, 12 rRNAs, 54 tRNAs, three other RNAs, and 187 pseudogenes. Only a single origin of replication (*oriC*) was found, at 1,863,144 bp, and no plasmids were detected (Supplementary Table 2). The lack of PacBio long reads overlapping the 5’ and 3’ genome regions indicated that the EP4 genome is linear.

EP4 grew on 1.0 mM 4-ethylphenol to stationary phase within 14 hours in shake flasks (Figure 1B). Growth on 4-ethylphenol was verified by plating CFUs (Figure 1C). GC-MS analysis indicated that 4-ethylphenol was completely removed from the medium during growth (Figure 1D), with no metabolites detected.

### Quasi-mapping-based quantification of prokaryotic gene expression

We used transcriptomics to identify 4-ethylphenol catabolic genes in EP4 without *a priori* bias. Transcriptome reads were aligned strand-wise to predict transcriptional start sites in the EP4 genome (Figure 2A). Read alignment is a common but time-consuming step in prokaryotic RNA-Seq pipelines (Supplementary Figure 1A). We therefore compared quasi-mapping to genomic coding regions using Salmon (Patro et al., 2017) with alignment-based read counting software. Salmon results were numerically (Supplementary Figure 1B) and statistically equivalent (*p_adj_* = 0.21) (Supplementary Figure 1C) to FeatureCounts, with strong correlation to RT-qPCR expression (R^2^_adj_ = 0.91, *p* < 0.001) (Supplementary Figure 1D). Salmon was about eight times faster than FeatureCounts, and superior to htseq in terms of total counts and accuracy, and was therefore used for gene quantification prior to differential expression analysis using *DESeq2*.

**Figure 2.**
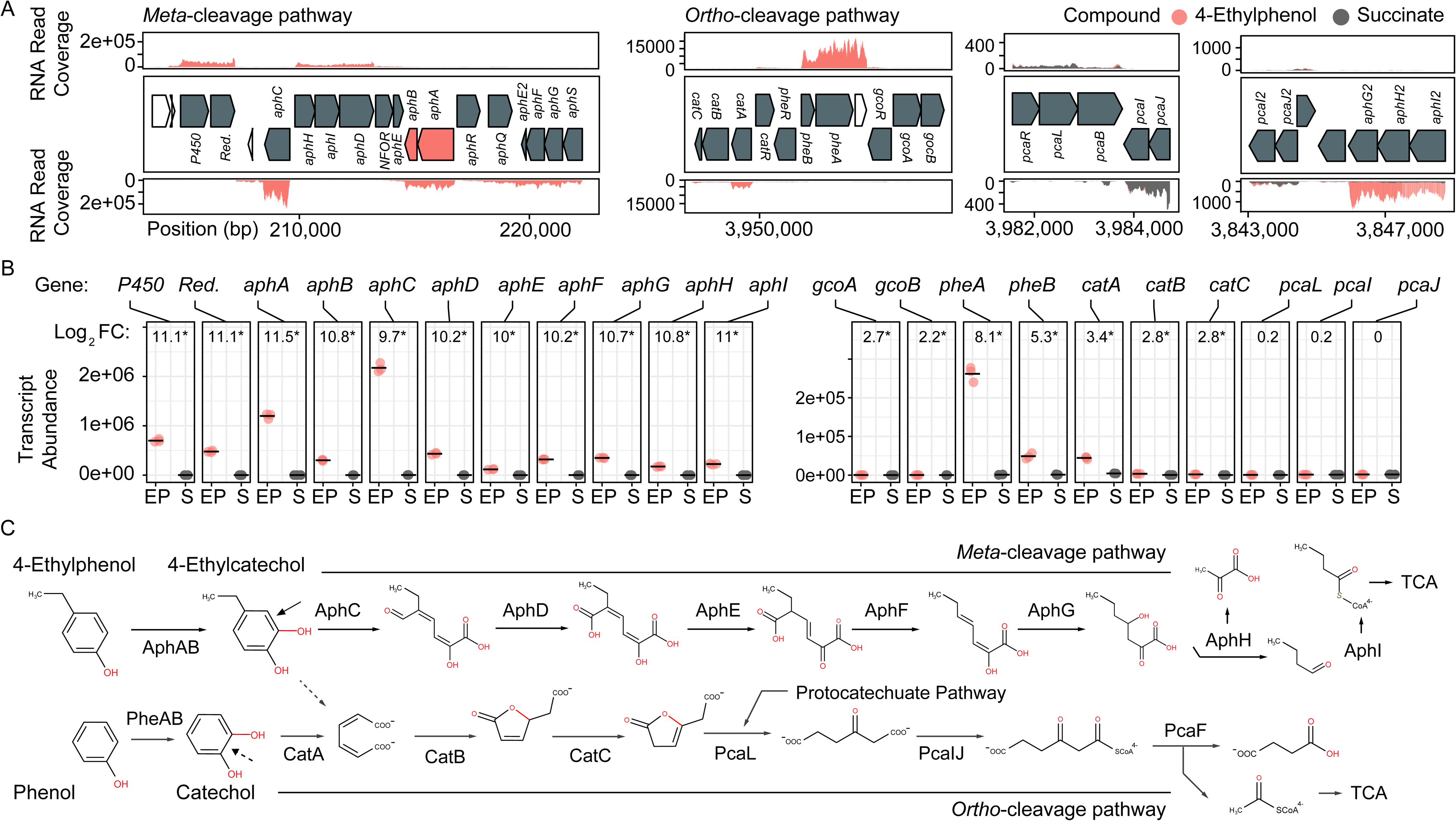
Transcriptomic and genomic identification of the 4-ethylphenol catabolic pathway genes in *Rhodococcus rhodochrous* EP4. A) RNA reads from cells grown on 4-ethylphenol or succinate mapped to the EP4 gene clusters encoding catechol meta-cleavage and ortho-cleavage. B) *Deseq2* differential-expression analysis showing log_2_ fold-change (FC) on 4-ethylphenol versus succinate (FDR-corrected p-values: * p_fdr_ < 0.001). Points show values for n = 3 replicates; horizontal bar indicates mean. P450, cytochrome P450 gene; Red., P450 reductase. C) Proposed funneling of 4-ethylphenol into the alkylcatechol *meta*-cleavage pathway (upper) and not the catechol *ortho*-cleavage pathway (lower). TCA, tricarboxylic acid cycle.

### Transcriptomic analysis of 4-ethylphenol metabolism via *meta*-cleavage

Growth on 4-ethylphenol versus succinate significantly modulated expression of 559 genes with *pfdr* < 0.001. Nine of the 16 most upregulated genes occurred in a cluster encoding a proposed alkylphenol catabolic pathway, *aphABCDEFGHIQRS* (Table 1). This cluster includes *aphAB*, encoding a two-component <u>a</u>lkyl<u>p</u>henol hydroxylase discussed below, and *aphC* (Figure 2B), encoding an extradiol dioxygenase that we subsequently identified as alkylcatechol 2,3-dioxygenase. The gene cluster is organized as four putative operons based on transcriptomic data and operon prediction with BPROM: *aphAB*, *aphHIDE*, *aphE2FGS*, *aphG2H2I2* (Figure 2A). The deduced Aph pathway catabolizes 4-ethylphenol to pyruvate and butyryl-CoA (Figure 2C), similar to the Dmp pathway of *Pseudomonas* sp. strain CF600 that catabolizes dimethylphenols (Shingler et al., 1992) and phenol (Powlowski and Shingler, 1994). Based on the transcriptomic data, the resultant butyryl-CoA is degraded to central metabolites by an aerobic fatty acid degradation pathway (Jimenez-Diaz et al., 2017) encoded by butyryl-CoA dehydrogenase genes (locus tags: C6369_RS06395, C6369_RS20140, C6369_RS07820, C6369_RS05465), enoyl-CoA hydratase (C6369_RS03325, C6369_RS19860), 3−hydroxybutyryl−CoA dehydrogenase (C6369_RS03325, C6369_RS06400) and acetyl-CoA acyltransferase (C6369_RS17095, C6369_RS15900, C6369_RS19850) (Supplementary Figure 2).

The *catABC* cluster encoding catechol 1,2-dioxygenase and other enzymes feeding into the β-ketoadipate pathway was also significantly upregulated during growth on 4-ethylphenol), although much less highly than the *aph* genes. No *ortho*-cleavage metabolites were detected in the culture supernatants (Figures 1D), and the genes encoding the downstream β-ketoadipate pathway, *pcaBLIJ*, were not up-regulated (Figure 2B). Overall, the data suggest that 4-ethyphenol is catabolized via *meta-*cleavage.

### Characterization of a two-component akylphenol hydroxylase, AphAB

We hypothesized that the highly upregulated *aphA* gene (L_2_FC = 11.5) encodes the oxygenase component of a novel alkylphenol monooxygenase, based on its location within the *aph* cluster as well as the phylogenetic and functional data presented below. The *aphB* gene, encoding a flavin reductase was co-transcribed with *aphA* (Figure 2B). The upregulation of *aphA* and *aphC* genes in EP4 during growth on 4-ethylphenol was confirmed using RT-qPCR (Supplementary Figure 3).

To establish the physiological role of AphAB from EP4, the oxygenase and reductase components were each overproduced in *E. coli* and purified to apparent homogeneity. The reconstituted AphAB_EP4_ hydroxylated 4-ethylphenol to 4-ethylcatechol (Figure 3A). The enzyme also catalyzed the hydroxylation of 4-methylphenol, 4-propylphenol (Figure 3B) and, to a much lesser extent, 4-NP (Figure 3C). However, AphAB _EP4_ did not detectably transform phenol (Figure 3B) or 4-HPA (Figure 3C). In an assay measuring cytochrome *c* reduction, AphB EP4 preferentially utilized NADH and flavin adenine dinucleotide (FAD) (Figure 3D), as reported for PheA2 (Gröning et al., 2014; Saa et al., 2010; Straube, 1987).

**Figure 3.**
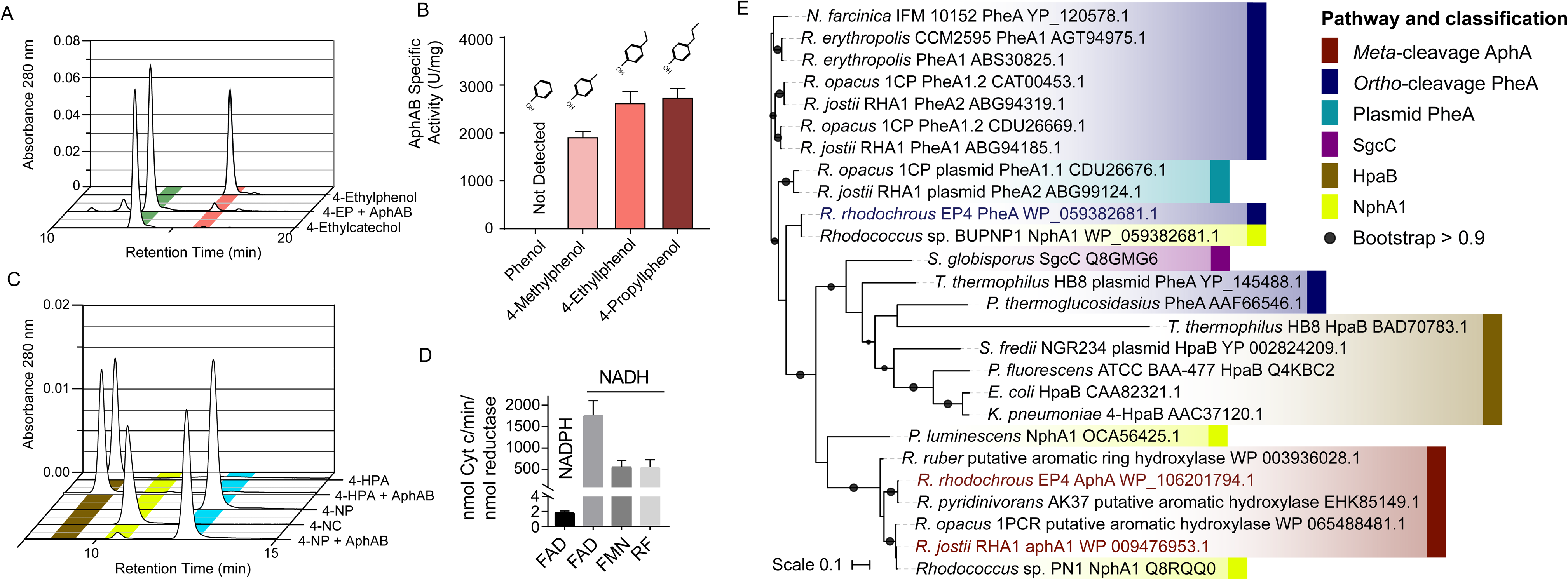
Characterization of AphAB_EP4_. A) Phylogenetic tree constructed using structural-based alignment and RAxML. B) Hydroxylation of 4-ethylphenol to 4-ethlycatechol by purified AphAB_EP4_. Reaction mixtures contained 20 µM of each enzyme component, 100 μM substrate, and were incubated overnight. C) Specific activity of AphAB_EP4_ for select phenols. Activity was measured using a coupled, spectrophotometric assay. D) Transformation of 4-HPA and 4-NP by AphAB_EP4_. Conditions as in C. E) Cofactor and substrate preference of AphB_EP4_. Reductase activity was measured using cytochrome *c*. HPA; hydroxyphenylacetate; NP, nitrophenol; FAD, flavin adenine dinucleotide; FMN, flavin mononucleotide; RF, riboflavin.

### Annotation of additional genes

During growth of EP4 on 4-ethylphenol, phenol hydroxylase genes, *pheA* and *pheB*, adjacent to the *cat* gene cluster, were additionally upregulated (Figure 2B, Supplementary Figure 3). In structure-based alignments, PheA_EP4_ and AphA_EP4_ clustered with separate 4-NP hydroxylases, rather than the clade of characterized phenol hydroxylases (Figure 3E). More specifically, PheA_EP4_ and AphA_EP4_ clustered most closely, respectively, with NphA1 from *Rhodococcus* sp. BUBNP1 (WP_059382681.1) (Sengupta et al., 2019) and NphA1 from *Rhodococcus* sp. PN1 (Q8RQQ0) (Takeo et al., 2003). Despite 100% sequence identity with NphA1_BUBNP1_, PheA _EP4_ was annotated based on sequence similarity to known phenol hydroxylases (Figure 3E). In support of this annotation, EP4 lacks a 4-NP catabolism gene cluster and was unable to grow on 4-NP, while it did grow on phenol (discussed below). PheA _EP4_ shares 82% identity with PheA1(1) (ABS30825.1) in *Rhodococcus erythropolis* UPV-1 (Saa et al., 2010) and 65% identity with a chlorophenol 4-monooxygenase (Q8GMG6) from *Streptomyces globisporus* (Liu et al., 2002) (Supplementary Table 4; Supplementary Figure 4). These similarities suggest that PheA_EP4_ may have broad substrate specificity.

In EP4, genes encoding AraC-family transcriptional regulators (TR) were found directly adjacent to and in the opposite orientation as *aphAB* and *pheAB* (Figure 2A) (Supplementary Figure 5). These AraC-family TRs were annotated as AphR and PheR, respectively. Another AraC-family TR is encoded by a gene immediately downstream of *aphR*, which has a distinct phylogeny from AphR (Supplementary Figure 5) and was annotated as AphQ. Finally, an IclR-family TR is encoded by the last gene of the *aphE2FGS* operon.

### Syntenic conservation of EP4 *aph* gene cluster in rhodococci

The above comparative analyses of hydroxylase proteins revealed homologs of the EP4 *aphA* gene in several other rhodococci. In RHA1, a putative *aphA* gene (Table 1) was previously annotated as an aromatic ring hydroxylase possibly involved in steroid degradation (McLeod et al., 2006) and there were three previously-identified *pheA* homologs (Supplementary Table 4) (Gröning et al., 2014). Local alignment of the 13 Aph proteins against proteins predicted from all 325 *Rhodococcus* genomes identified 75 strains with full or partial (≥7 genes) putative Aph pathways, including RHA1 (Supplementary Figure 5A). Related pathways were also found in other Actinobacteria, but this study focused on the rhodococcal pathway. RHA1 *aph* genes displayed syntenic conservation with the EP4 *aph* cluster (Figure 4), except for an additional butyryl-CoA dehydrogenase gene (RHA1_RS18815). The *aphCHIDE* region was conserved in all *aph*-containing genomes based on *nucmer* alignment (Figure 4, Supplementary Figure 5B).

**Figure 4.**
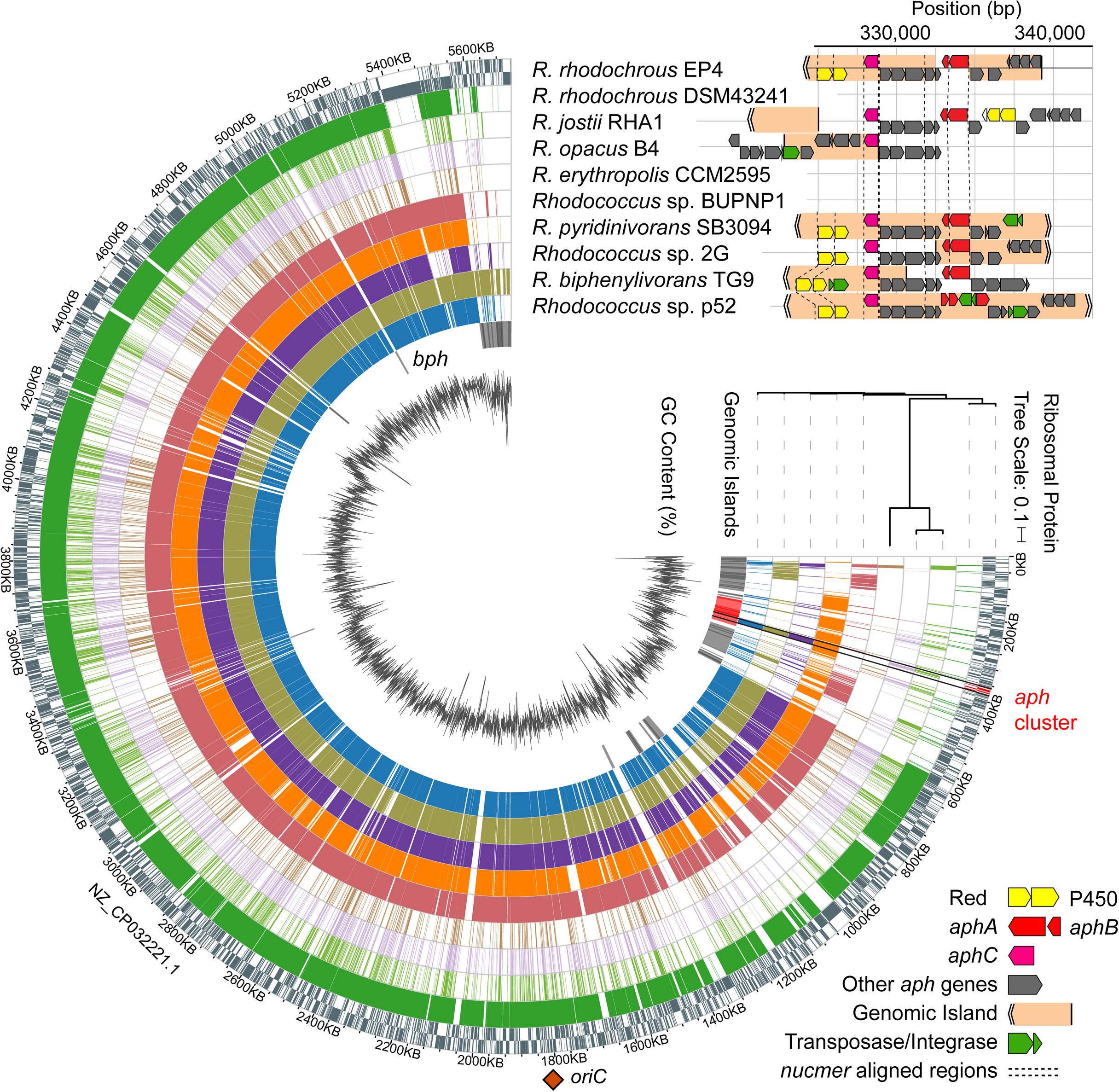
Identification of a putative *aph* genomic island in rhodococci. A) Alignment of *Rhodococcus* genomes to EP4 reference with *nucmer* ordered by RAxML tree calculated from concatenated alignment of 16 ribosomal protein sequences; predicted RHA1 genomic islands; GC content (%); and syntentic organization of *aph* gene clusters showing genomic islands predicted using IslandViewer4. *Nucmer* alignment regions shown with dashed line. *aphA*, 4-alkylphenol 3-monooxygenase, oxygenase gene; *aphB*, 4-alkylphenol 3-monooxygenase, reductase gene; *aphC*, alkylcatechol 2,3-dioxygenase gene; *bph*, biphenyl catabolism gene cluster; P450, cytochrome P450 gene; Red., P450 reductase. The single origin of replication (*oriC*) shown with orange diamond. Detail of the *aph* region alignments in Supplementary Figure 6B.

The EP4 and RHA1 Aph proteins shared 66.7% to 90.4% identity (median = 85.7%). Consistent with the occurrence of the putative Aph pathway in RHA1, this strain grew on 4-ethylphenol (Figure 5A), with concomitant upregulation of the *aph* genes (Figure 5B). Specifically, *aphA* and *aphC*) were highly upregulated (L_2_FC, 14.3 and 10.0, respectively). In contrast to EP4, when RHA1 grew on 4-ethylphenol, it did not upregulate any of its three *pheA* genes or any of the ring-cleavage dioxygenase genes associated with the *pheA* genes, including *catA* (RHA1_RS11595), *catA2* (RHA1_RS35920) and a plasmid-borne catechol 2,3-dioxygenase gene (RHA1_RS35970) (McLeod et al., 2006) (Figure 5B).

**Figure 5.**
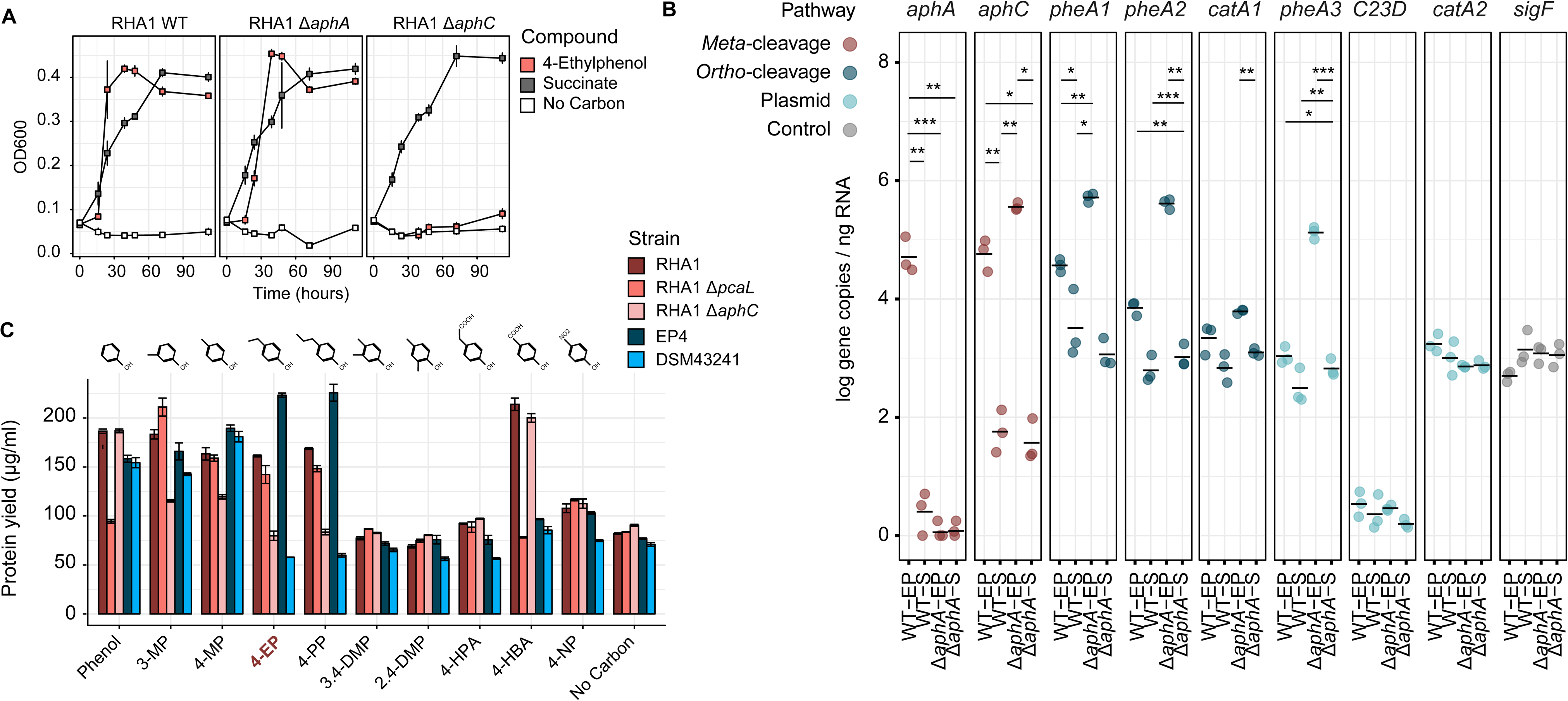
Molecular genetic analysis of 4-ethylphenol catabolism in RHA1. A) Growth of WT, Δ*aphA* and Δ*aphC1* RHA1 strains on 1 mM 4-ethylphenol or 2 mM succinate. B) Protein yield of WT, Δ*aphC*, and Δ*pcaL* RHA1 strains as well as EP4 and *R*. *rhodochrous* DSM43241 on phenolic substrates. Protein measured after 24 hours incubation. C) Expression of select genes in WT and Δ*aphA* RHA1 strains during growth on 1 mM 4-ethylphenol or 2 mM succinate using RT-qPCR. Colors indicate cleavage pathway. Points and horizontal bars show individual measurements (n = 3) and mean. Significance levels following Bonferroni-corrected two-tailed Student’s t-tests (*, p_bon_<0.05; **, p_bon_<0.01; ***, p_bon_<0.001). DMP, dimethylphenol; EP, ethylphenol; HBA, hydroxybenzoic acid; HPA; hydroxyphenylacetate; MP, methylphenol; NP, nitrophenol; PP, propylphenol.

### Gene deletion analysis of 4-alkylphenol ring cleavage

Because RHA1 is genetically-tractable, we constructed Δ*aphA* and Δ*aphC* deletion mutants and used these together with an existing Δ*pcaL* mutant to further investigate 4-ethylphenol catabolism. The Δ*aphC* mutant did not grow on either 1.0 mM 4-ethylphenol or 4-propylphenol (Figure 5AB) demonstrating that both compounds are exclusively metabolised by *meta-*cleavage. However, Δ*aphA* did grow on 4-ethylphenol (Figure 5A). It appears that one or more of three PheA homologs from RHA1 may catalyze 4-ethylphenol hydroxylation and compensate for the deletion of *aphA*. While the corresponding *pheA* genes were not upregulated in wild-type RHA1 growing on 4-ethylphenol versus succinate, they were upregulated 7.6 to 8.8 L_2_FC in the Δ*aphA* mutant (Figure 5B), while the plasmid-borne *C23D* gene was not upregulated. Finally, the Δ*pcaL* mutant grew on alkylphenols but did not grow on either phenol or 4-HBA, indicating that the latter two substrates are catabolized solely via *ortho-*cleavage pathways (Figure 5C).

### Identification of a putative *aph* genomic island

The *aph* gene cluster (approximately 17 kb) occurs within 117 kb and 4.2 kb regions predicted to be two of 61 GIs (or 38 non-overlapping GI regions) identified in EP4 using IslandViewer4 (Figure 4; Supplementary Table 2). These GI elements do not include *aphAB* or *aphE* in EP4, but do in three other *Rhodococcus* strains. These putative GIs are located near the 3’ end of the EP4 chromosome assembly in a 600-kb region of apparent genomic instability, as it contains high insertion sequence (IS) density (Figure 1A) and 36 predicted GIs (Figure 4). The GI-like characteristics of these elements containing the *aph* cluster include a −5.8% deviation from the mean GC content, presence of mobility genes (integrases, transposases, insertion sequences) and absence of the region in closely-related genomes following alignment (Langille et al., 2008) (Figure 4; Supplementary Figure 6). They are not located near a tRNA sequence, indicating that it is not likely a mobile integrative and conjugative element (ICE) (Burrus and Waldor, 2004). More generally, 7.4% of the EP4 genome and 9.2% of the RHA1 genome were predicted to occur on GIs (Supplementary Table 2).

Analysis of 37 complete, full-length *Rhodococcus* genomes found that 16 carried genes encoding a complete Aph pathway. The *aph* genes are predicted to be fully or partially contained in a GI in six of these strains and to occur immediately downstream of a GI in a seventh, RHA1. This genomic region was conserved in three *Rhodococcus* clades: one containing EP4 and *R. pyridinivorans* strains, one containing *R. jostii*, *R. opacus* and *Rhodococcus wratislaviensis* strains and one containing *Rhodococcus* sp. Eu32 (Supplementary Figure 6). With the exception of a partial *aph* cluster in *R. rhodochrous* ATCC 21198, the *aph* genes were not found in any of the 13 other *R. rhodochrous* genomes including strain DSM43241. Accordingly, DSM43241 could not grow on 4-ethylphenol and 4-propylphenol, but grew on phenol, 4-HBA, 3-methylphenol and 4-methylphenol (Figure 5C).

## Discussion

In this study, we used a variety of approaches to identify an Aph pathway responsible for the catabolism of alkylphenols via *meta*-cleavage in *Rhodococcus*. Catabolism is initiated by AphAB, a two-component hydroxylase that transforms the alkylphenol to the corresponding alkylcatechol (Figures 2,3). To date, only six-component proteobacterial alkylphenol hydroxylases have been reported (Arai et al., 2000; Jeong et al., 2003). The ensuing Aph pathway generates pyruvate and an acyl-CoA following *meta*-cleavage of 4-alkylcatechol. The length of the acyl-CoA produced depends on the alkyl side chain of the growth substrate. This is in contrast to the Phe and Nph pathways which catabolize phenol and 4-NP, respectively, via *ortho*-cleavage (Sengupta et al., 2019; Szőköl et al., 2014; Takeo et al., 2008).

The activity of AphAB_EP4_ is consistent with its phylogenetic relationship with two-component phenolic hydroxylases. Thus, the clade containing AphA_EP4_ and AphA_RHA1_ includes an NphA1 but no characterized PheA or HpaB (Figure 3). In keeping with this classification, AphAB_EP4_ had weak activity with 4-NP (Figure 3C) but did not detectably transform phenol or 4-HPA. However, the determinants of substrate specificity of these enzymes are not clear. The catalytic residues of these hydroxylases (Chang et al., 2016; Kim et al., 2007) are conserved in AphA: Arg119, Tyr123 and His161 (AphA_EP4_ numbering; Supplementary Figure 4). In a structurally characterized HpaB:4-HPA binary complex, the substrate’s carboxylate is coordinated by Ser197 and Thr198 (Kim et al., 2007). In PheA, NphA and AphA, these residues are His214 and Tyr215, suggesting that they do not contribute to the enzyme’s substrate specificity despite their predicted interaction with the *para*-substituent of the substrate.

The Aph pathway is similar to the Dmp pathway described in *Pseudomonas* sp. strain CF600 (Shingler et al., 1992). However, it is clear that the Aph pathway has a distinct substrate specificity because neither EP4 nor RHA1 grew on 2,4- or 3,4-DMP (Figure 4E) and *aph* pathway mutants grew on phenol. We had previously suggested that some of the *aph* genes could be involved in steroid degradation (McLeod et al., 2006) due to their similarity to known steroid catabolic genes (Van der Geize et al., 2007). Further, in a recently published structure of AphC_RHA1_, the enzyme was identified as 2,3-dihydroxybiphenyl dioxygenase (Table 1). However, AphC is encoded in a gene cluster upregulated on 4-alkylphenols and is essential for growth of RHA1 on those compounds, supporting annotation of this rhodococcal Aph *meta-*cleavage pathway, with AphC as an alkylcatechol 2,3-dioxygenase.

4-Ethylphenol strongly induced *aphAB* expression. This is likely due to positive induction of the AphR TR, just as phenol activates *pheA2A1* expression by PheR in RHA1 (Szőköl et al., 2014), and 4-NP activates *npaA2A1* expression by NphR in *Rhodococcus* sp. PN1 (Takeo et al., 2008). AphR, PheR and NphR are all AraC-family TRs. AphR and PheR may play a role in the unexpected ability of the RHA1 Δ*aphA* mutant to grow on 4-ethylphenol. The lack of *pheA1-3* expression in wild-type RHA1 (Figure 5B) strikingly contrasts with the upregulation of these genes in the Δ*aphA* mutant (Figure 5C). Binding regions for PheR_RHA1_ and predicted AphR_RHA1_ binding sites overlap by 10/18 nt (Supplementary Figure 5B), suggesting the potential for non-target regulation, as Szőköl et al. (2014) argue for *R. erythropolis* CSM2595 PheR and XylS. This overlap is much less in predicted PheR_EP4_ and AphR_EP4_ binding regions (3/18 nt). Indeed, the *aphA* promotor (−10: CAGGAG; −35: CCGTCT) (Supplementary Figure 5C) bears more similarity to the T80 promotor of *Mycobacterium tuberculosis* (Bashyam et al., 1996) than with the rhodococcal *pheA* promotors. It is also noted that two of the *pheA* genes in RHA1 are subject to catabolite repression (Szőköl et al., 2014). It is possible that this repression is relieved in the Δ*aphA* mutant that can’t metabolise 4-ethylphenol. Related to this, homologs of PheAB in *R. opacus* 1CP hydroxylated 4-methylphenol with about twice the specific activity as with phenol (Gröning et al., 2014), further suggesting that the RHA1 PheABs may hydroxylate 4-ethylphenol.

In addition to 4-ethylphenol, alkylguaiacols and alkylsyringols commonly occur in lignin depolymerization streams (Anderson et al., 2016; Asawaworarit et al., 2019; Guo et al., 2017; Jiang et al., 2014; Kim et al., 2015; Ye et al., 2012). Interestingly, genes encoding a cytochrome P450 and reductase are linked to the *aph* clusters in some rhodococci (Figures 2,4). Further, these were the second and third most highly upregulated genes in EP4 during growth on 4-20 ethylphenol versus succinate (both L_2_FC = 11.1) (Figure 2B). The P450 shares 65% sequence identity with a guaiacol *O*-demethylase (Mallinson et al., 2018), suggesting that the rhodococcal enzyme has a similar role, and that these strains may also funnel methoxylated compounds into the Aph pathway.

The ability of RHA1 and EP4 to catabolize 4-ethylphenol and other alkylphenols is of potential use in upgrading lignin streams generated by RCF and other depolymerization strategies. The Aph *meta*-cleavage pathway harboured by these strains contrasts with the *ortho*-cleavage pathways targeted to date in the design of biocatalysts for lignin valorization (Abdelaziz et al., 2016; Barton et al., 2018; Beckham et al., 2016). This is largely due to the identified economic potential of some of the *ortho*-cleavage metabolites. For example, *cis*,*cis*-muconate resulting from *ortho*-cleavage can be used to make adipic acid and terephthalic acid (Barton et al., 2018; Beckham et al., 2016; Xie et al., 2014). However, alkylphenols may be funneled through *ortho*-cleavage by oxidizing the *para*-side chain to 4-hydroxyacetophenone or hydroquinone (Darby et al., 1987). Alternatively, oleaginous *Rhodococcus* strains such as RHA1 may be modified to use the Aph pathway to produce lipid-based commodity chemicals (e.g., Round et al., 2017) offering a method for valorization of alkylphenols via fatty-acid synthesis in this genus.

We found genes encoding the Aph pathway in several *Rhodococcus* strains, including oleaginous strains such as RHA1 and *R. opacus* B4 (Figure 3A, Figure 4). However, the absence of an *aph* cluster in most *R*. *rhodochrous* strains (e.g., DSM43241) demonstrates that phylogenetically-related strains can have important metabolic differences. Previous studies suggested that recent horizontal gene transfer did not play a large role in generating RHA1’s considerable catabolic capabilities (McLeod et al., 2006). In several rhodococci, putative GIs did not contain all of the *aph* genes. This could represent the imprecision of the prediction tools, incomplete amelioration of the element, or reflect that these GIs arose from separate insertion events. Patchwork *aph* GIs are consistent with the theorized modular origins of GIs (Juhas et al., 2009). Fermentation of plant-derived aromatic compounds, including cinnamic acids, by yeasts and lactic acid bacteria can naturally produce alkylphenols (Caboni et al., 2007; Kridelbaugh et al., 2010). The apparent complete loss of the *aph* genes in other *R. rhodochrous* strains may result without selective alkylphenol exposure if it otherwise has a deleterious effect on overall fitness. Testing these regions for their excision capacity was beyond the scope of this work, but remains an intriguing prospect. We posit that the presence of aromatic compounds in compost selected for microorganisms capable of 4-ethylphenol catabolism, including EP4.

In this study we described a newly-isolated, 4-ethylphenol-catabolizing strain, EP4, a novel alkylphenol hydroxlase, AphAB, and its role in funneling alkylphenols into the Aph *meta*-cleavage pathway in some *Rhodococcus* strains. We showed that this pathway is associated with putative GIs, primarily found in strains from contaminated soil environments. Characterizing 4-ethylphenol metabolism in EP4 and RHA1 advances our capacity for bio-refinement of reductively depolymerized lignin subunits from sustainable chemical feedstocks.

## Acknowledgements

This study was supported by a research contract from Genome BC (SIP004) and a grant from the Natural Sciences and Engineering Research Council of Canada (STPGP 506595-17). All sequencing was performed at the McGill University and Génome Québec Innovation Centre (Montreal, CAN). We thank Andrew Wilson and Alexandra Booth for their assistance with laboratory experiments.

## Supplementary Figure Captions

Supplementary Figure 1. Quantitative mapping of prokaryotic RNASeq reads. A) Compute time in seconds for read mapping steps for three methods of feature quantification: FeatureCounts (FC), HTSeq and Salmon. Timing benchmarks calculated on a 256 Gb, 32-core Ubuntu 14.04.5 server, running eight cores per analysis. B) Total counts produced by each method. C) PCoA ordination of read counts of all genomic features with results of pairwise PERMANOVA. D) Linear regression of RNASeq method against RT-qPCR transcript abundance showing R2adj. All *p*-values < 0.0001). 4-EP, 4-ethylphenol; 4-PG, 4-propylguaiacol; S, succinate. Full 4-PG results can be found in a separate manuscript (Fetherolf et al., unpublished).

Supplementary Figure 2. Transcriptomic analysis of the metabolism of butyryl-CoA produced from 4-ethylphenol catabolism by the Aph pathway in EP4. The top panel shows the predicted pathway of metabolism showing the production of the acyl-CoA during the final step of the Aph pathway. The bottom panels show plots of −log_10_ transformed *p*-values plotted against log_2_ fold change for all putative homologs of the genes encoding each step of acyl-CoA metabolism during growth on 1.0 mM 4-ethylphenol versus succinate calculated using *DeSeq2*.

Supplementary Figure 3. RT-qPCR expression of *aphA*, *aphC*, *pheA*, *catA* and *sigF* (control) genes during growth of EP4 on 1 mM 4-ethylphenol (EP), 2 mM succinate (S) and 1 mM benzoate. N = 3, log10 gene copies ng^−1^ total RNA). Table shows log_2_ fold change and *p*-value following Bonferroni-corrected two-tailed Student’s t-tests: *, *p_bon_* < 0.05, **, *p_bon_* < 0.01.

Supplementary Figure 4. TCoffee-Expresso-based structure alignment of EP4 AphA and select homolog amino acid sequences. See Figure 3 for sequence identification.

Supplementary Figure 5. Investigating the role of AphR in *aphAB* expression. A) RAxML-tree from TCoffee-aligned AraC-family transcriptional regulatory proteins. Red indicates previously identified proteins. B) Alignment of EP4 and RHA *aphA*-AphR and *pheA*-PheR binding regions; inset: organization of EP4 and RHA1 *aphBAR* and *pheRBA* genes. D) Map of the *aphR*-*aphA* intergenic region showing the AphR binding site predicted from alignment with CCM2595 PheR binding regions (yellow, dotted underline), *aphA* promotors (solid underline), *aphA* transcriptional start site (TSS) (purple) and predicted coding start (CS) (pink). Strains: ATCC39116, *Amycolatopsis* sp. ATCC39116; BUPNP1, *Rhodococcus* sp. BUPNP1; CCM2595, Rhodococcus *erythropolis* CCM2595; KL28, *Pseudomonas* sp. KL28.

Supplementary Figure 6. The *aph* cluster in *Rhodococcus* genomes. A) RAxML phylogenetic tree calculated from MUSCLE alignment of concatenated sequences for ribosomal proteins L2, L3, L4, L5, L6, L14, L16, L18, L22, L24, S3, S8, S10, S17 and S19 from 325 *Rhodococcus* ssp. genomes showing genomes for which BLASTp evalues were < 10^−100^ for at least 7 of the 13 EP4 Aph amino acid sequences (gold boxes). Also showing the *Rhodococcus opacus*/*jostii/wratislaviensis* clade (green), *Rhodococcus rhodochrous*/*pyradinivorans* clade (gold), and *Rhodococcus* sp. Eu-32 (peach). B) Detail from Figure 4 showing *nucmer* alignment of select *Rhodococcus* genomes to the EP4 *aph* gene cluster including a portion of the 117-kb *aph* gene-containing genomic island predicted using IslandViewer4 (grey), showing *nucmer* aligned regions (black dashed lines), median GC content (red dashed line) and GC content in 200 bp segments (solid grey line).

